# Optimization of energy production and central carbon metabolism in a non-respiring eukaryote

**DOI:** 10.1101/2022.12.29.522219

**Authors:** Sara Alam, Ying Gu, Polina Reichert, Jürg Bähler, Snezhana Oliferenko

**Affiliations:** The Francis Crick Institute, 1 Midland Road, London, NW1 1AT, UK; Randall Centre for Cell and Molecular Biophysics, School of Basic and Medical Biosciences, King’s College London, London, SE1 1UL, UK; Medical Research Council London Institute of Medical Sciences, Du Cane Road, London, W12 0NN, UK; School of Biological and Behavioural Sciences, Queen Mary University of London, Mile End Road, London E1 4NS, UK; Institute of Healthy Ageing, Department of Genetics, Evolution and Environment, University College London, London, WC1E 6BT, UK

## Abstract

Most eukaryotes respire oxygen, using it to generate biomass and energy. Yet, a few organisms lost the capacity to respire. Understanding how they manage biomass and energy production may illuminate the critical points at which respiration feeds into central carbon metabolism and explain possible routes to its optimization. Here we use two related fission yeasts, *Schizosaccharomyces pombe* and *Schizosaccharomyces japonicus*, as a comparative model system. We show that although *S. japonicus* does not respire oxygen, unlike *S. pombe*, it is capable of efficient NADH oxidation, amino acid synthesis and ATP generation. We probe possible optimization strategies using stable isotope tracing metabolomics, mass isotopologue distribution analysis, genetics, and physiological experiments. *S. japonicus* appears to have optimized cytosolic NADH oxidation via glycerol-3-phosphate synthesis. It runs a fully bifurcated TCA ‘cycle’, supporting higher amino acid production. Finally, it uses the pentose phosphate pathway both to support faster biomass generation and as a shunt to optimize glycolytic flux, thus producing more ATP than the respiro-fermenting *S. pombe*. By comparing two related organisms with vastly different metabolic strategies, our work highlights the versatility and plasticity of central carbon metabolism in eukaryotes, illuminating critical adaptations supporting the preferential use of glycolysis over oxidative phosphorylation.

## Introduction

Carbon metabolism, producing biomass and energy, is essential for growth and survival in most organisms. Establishing the rules of carbon metabolism is critical for our understanding of life, from evolution to development to disease [1-6]. In glycolysis, a molecule of glucose is catabolized to pyruvate, generating two ATP molecules. Pyruvate may be decarboxylated to acetaldehyde and then reduced to ethanol through fermentation, which oxidises the NADH generated by glycolysis rendering this metabolic strategy redox-neutral. Alternatively, in respiration, pyruvate may be converted to acetyl-CoA which then enters the tricarboxylic acid (TCA) cycle. Each round of the TCA cycle provides precursors for amino acids and nucleotides, as well as generating NADH and succinate. In turn, NADH and succinate are oxidised via the electron transport chain (ETC), generating a potential across the inner mitochondrial membrane to power ATP synthesis. In yeasts that do not have the proton-pumping ETC complex I, respiration together with the initial catabolism of glucose via glycolysis can generate up to 16-18 ATP per glucose [7-11]. Most eukaryotes are capable of both respiration and fermentation; cells choose one metabolic strategy over the other depending on, for example, the nutritional environment [12, 13]. Indeed, Crabtree-positive model yeasts such as *Schizosaccharomyces pombe* (*S. pombe*) and *Saccharomyces cerevisiae* (*S. cerevisiae*) channel more glucose towards fermentation when ample glucose is available [13, 14]. While fermentation is less efficient at generating ATP, it produces it quickly and at a low cost, while providing Crabtree-positive species with competitive advantages in the wild, such as diverting glucose away from other organisms [1, 11, 15-17].

In principle, whereas respiration may be repressed in glucose-rich environments, ETC activity and oxidative phosphorylation (OXPHOS) can be of value to growing cells. Growth of non-respiring *S. cerevisiae* and *S. pombe* is improved by amino acid supplementation, suggesting that biomass production is limited when respiration is blocked [13, 18, 19]. Additionally, respiration may be the most efficient method not only for ATP synthesis but also for NADH oxidation that is essential to support growth [20, 21]. How is eukaryotic central carbon metabolism structured to overcome the limitations associated with the loss of respiration?

The fission yeast *Schizosaccharomyces japonicus* (*S. japonicus*), together with the more widely used relative *S. pombe*, provide an attractive composite model system to study this question. Unlike *S. pombe, S. japonicus* thrives both in the presence and the absence of oxygen [22-27]. Despite encoding most genes required for respiration, it does not produce coenzyme Q, does not grow on a non-fermentable carbon source glycerol, and does not consume oxygen during vegetative growth in glucose, unlike its sister species [22, 23, 26, 28]. To our knowledge, such traits have not been demonstrated for other tractable unicellular eukaryotes, positioning *S. japonicus* as a unique model system to study non-respiratory metabolism.

The two fission yeast species have already furthered our understanding of the evolution of cell biological processes, such as mitotic nuclear envelope remodelling, cell polarity, cytokinesis and membrane organisation [29-36]. We reasoned that understanding how *S. japonicus* manages its fully fermentative lifestyle might provide fundamental insights into the wiring of central carbon metabolism.

Here, we establish *S. japonicus* as a powerful model system for understanding the physiology of non-respiring cells. We further explore the impact of this metabolic strategy on NADH oxidation, biomass production and ATP synthesis. Our results suggest that *S. japonicus* is surprisingly well-adapted to grow without respiring oxygen, in stark contrast to its respiro-fermenting sister species *S. pombe*.

## Results

### *S. japonicus* does not respire oxygen in a variety of physiological situations

The *cox6* gene encodes an evolutionarily conserved subunit of the ETC complex IV, which is essential for respiration in *S. pombe* [37]. To test possible contribution of respiration to *S. japonicus* physiology, we analysed the growth requirements and oxygen consumption in the wild type and *cox6Δ S. japonicus* and the corresponding strains of *S. pombe*.

Consistent with previously published results [26], in rich glucose medium the wild type *S. japonicus* exhibited a much lower oxygen consumption rate than *S. pombe* (Fig. 1A). Whereas the deletion of *cox6* critically decreased oxygen consumption in *S. pombe*, we observed no such effect in *S. japonicus* (Fig. 1A). This result suggested that the minimal oxygen consumption in vegetatively growing wild type *S. japonicus* and *cox6Δ S. pombe* cells was likely due to non-respiratory oxygen-consuming biochemical processes [38-40]. *S. japonicus* did not grow on the non-fermentable carbon sources, glycerol and galactose (Fig. S1A), indicating that the lack of oxygen consumption in glucose was not due to respiratory repression. This behaviour was observed in several wild isolates [26, 41] and in *S. japonicus var. versatilis* [42, 43], suggesting that this feature is a species-wide trait (Fig. S1B).

**Figure 1.**
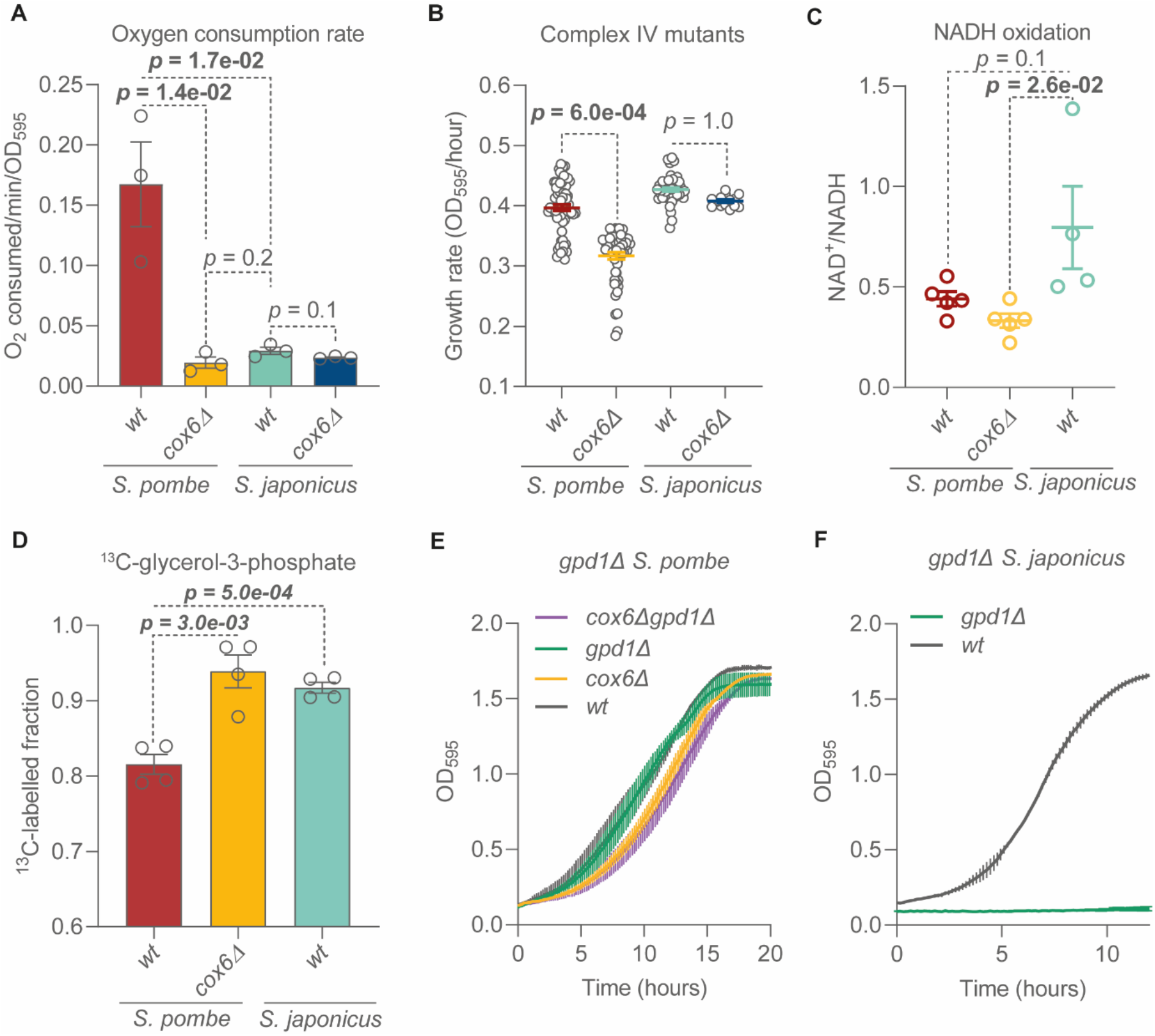
*S. japonicus* does not respire but oxidizes NADH efficiently. (**A**) Oxygen consumption rates of *S. pombe* and *S. japonicus* wild type (wt) and *cox6Δ* cultures in rich YES medium. Mean values are derived from three biological replicates. (**B**) Growth rates of *S. pombe* and *S. japonicus* wild type (wt) and *cox6Δ* cells in minimal EMM medium. Mean values are derived from at least five biological repeats with three technical replicates each. (**C**) Total cellular NAD^+^/NADH ratios of *S. pombe* and *S. japonicus*. Mean values are derived from at least four biological replicates. (**D**) Proportion of ^13^C-labelled glycerol-3-phosphate in *S. pombe* and *S. japonicus* cultures after 60 minutes of exposure to ^13^C_6_-glucose in minimal EMM medium. Mean values represent two biological and two technical replicates. (**A**-**D**) Error bars represent ±SEM; *p* values are derived from unpaired t-test. (**E, F**) Growth curves of *S. pombe* and *S. japonicus* cultures of indicated genotypes in YES. Error bars represent ±SD. Shown are the mean values of OD_595_ readings derived from three technical replicates, representative of three independent biological experiments.

*S. pombe* relies on ETC activity for efficient growth in minimal media, when cells are forced to synthesize most biomass precursors [13, 18]. In contrast, the deletion of *cox6* in *S. japonicus* did not impact the mean growth rate in Edinburgh minimal medium (EMM), unlike in its sister species (Fig. 1B). We tested whether other physiological processes in *S. japonicus* could be affected by the genetic disruption of the ETC. We did not observe any defects in yeast-to-hyphae transition and hyphal growth in *S. japonicus cox6* mutants (Fig. S1C). Respiration is essential for sporulation in budding yeast [44] and plays a significant role in *S. pombe* mating and sporulation [45]. However, *cox6Δ S. japonicus* mutants showed normal mating and sporulation efficiency (Fig. S1D). Thus, the loss of ETC function largely did not affect key physiological states of *S. japonicus*, at least in our laboratory conditions.

A key function of the ETC is the oxidation of NADH [20]. To test whether NADH oxidation is impacted by the evolutionary loss of respiration in *S. japonicus*, we measured the whole-cell NAD^+^/NADH ratios in both sister species. *S. japonicus* exhibited a similar NAD^+^/NADH ratio to the wild type *S. pombe*; in fact, this ratio was higher than in non-respiring *S. pombe* mutants (Fig. 1C). This finding suggests that *S. japonicus* has evolved ETC-independent mechanisms to efficiently oxidise NADH.

Anaerobically grown *S. cerevisiae* depends on the reduction of dihydroxyacetone phosphate (DHAP) to glycerol-3-phosphate (G3P) by the glycerol-3-phosphate dehydrogenase Gpd1 to oxidise cytosolic NADH [46-48]. We hypothesized that *S. japonicus* may rely on this reaction to sustain fermentative growth. We used stable isotope tracing coupled to gas chromatography mass spectrometry (GS-MS) to explore the rate of incorporation of glucose-derived carbons into G3P [49, 50]. After 60 minutes of exposure to ^13^C_6_-glucose, both *S. japonicus* and the non-respiring *S. pombe cox6Δ* mutants showed a significantly higher rate of ^13^C incorporation into the G3P pool, as compared to the wild type respiro-fermenting *S. pombe* (Fig. 1D). This result suggests that both fission yeasts might make use of DHAP reduction to sustain NADH oxidation in the absence of respiration. Strikingly, while the deletion of *gpd1* in *S. pombe* did not affect its growth, regardless of respiratory activity (Fig. 1E), *S. japonicus gpd1Δ* cells were virtually incapable of growth even in the rich YES medium, and could not be maintained at all in the minimal EMM (Fig. 1F). We conclude that *S. japonicus* critically depends on Gpd1 activity, whereas *S. pombe* may have additional mechanisms to oxidise cytosolic NADH.

### *S. japonicus* operates an obligately bifurcated TCA ‘cycle’ with endogenous bicarbonate recycling, whereas the respiro-fermenting *S. pombe* maintains both bifurcated and cyclic TCA variants

Respiration is typically associated with the TCA cycle, which allows for the additional oxidation of glucose, enabling more ATP production [10]. In addition to its role in supporting respiration, the TCA cycle generates crucial biomass precursors, alpha-ketoglutarate (aKG) and oxaloacetate (OAA), which are used to synthesize glutamate, aspartate, and other amino acids and metabolites (Fig. 2A). In anaerobic *S. cerevisiae* cultures, the TCA cycle has been shown to bifurcate, with an oxidative branch running from acetyl-CoA to aKG, and a reductive branch starting from the carboxylation of pyruvate to oxaloacetate, with subsequent conversion to malate, fumarate and succinate [51]. In the absence of respiration, bifurcation of TCA reactions presumably enables cells to support OAA and aKG synthesis in a redox-neutral manner (Fig. 2B). The TCA cycle architecture has not yet been probed in *S. pombe* and *S. japonicus*. We assessed it using stable isotope tracing metabolomics, quantifying the ratios of different TCA-intermediate isotopologues after feeding cells with ^13^C_6_-glucose for 30 minutes (Fig. 2A, B). To mimic the ‘broken’ TCA cycle, we included the *S. pombe* mutant lacking Sdh3, the key subunit of the succinate dehydrogenase (SDH) complex [52-54]. We used fumarate, the product of SDH, as a diagnostic metabolite.

**Figure 2.**
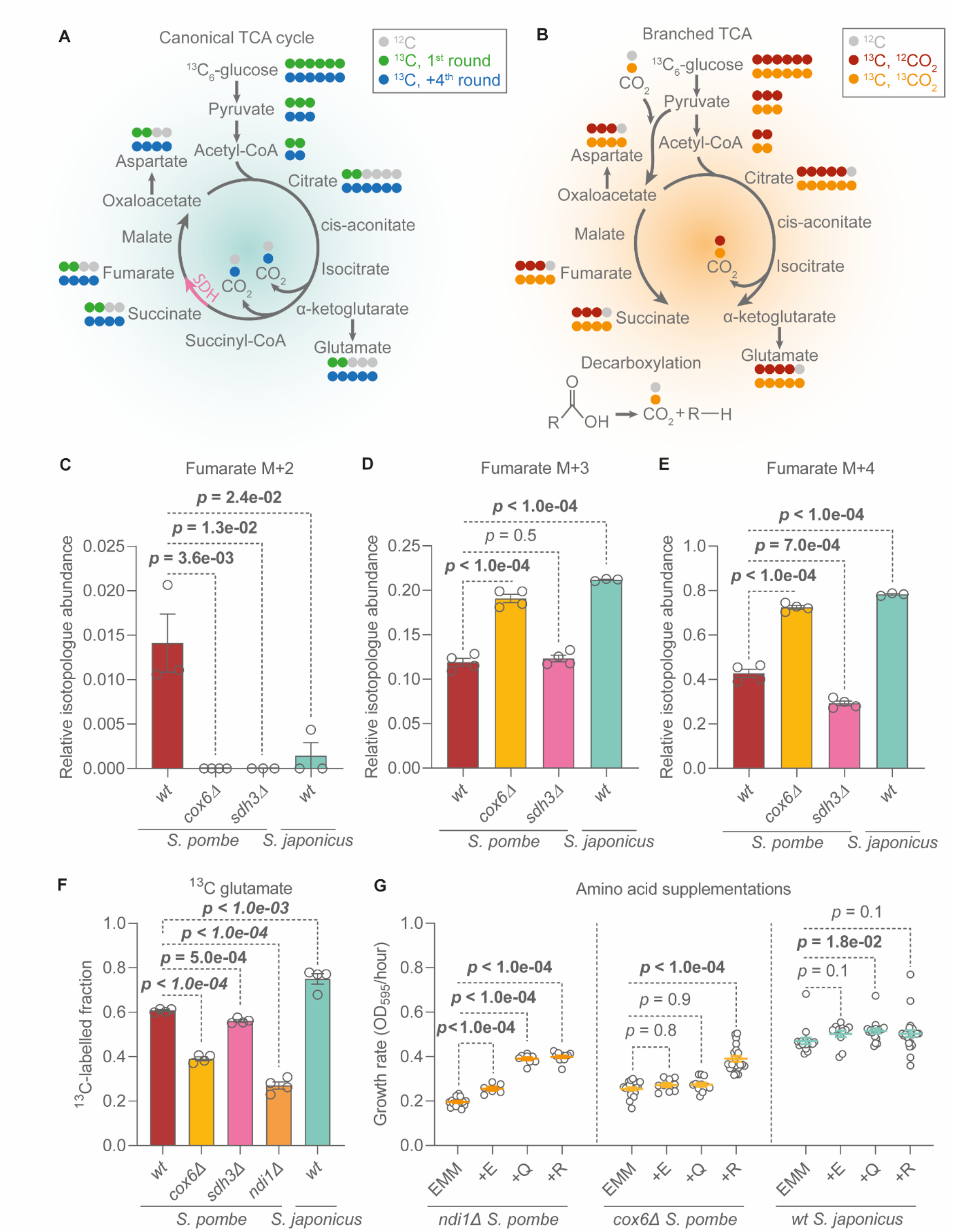
*S. japonicus* operates a bifurcated TCA ‘cycle’ and efficiently synthesizes TCA-derived amino acids. (**A**) Illustration of isotopologues of TCA intermediates expected when the oxidative TCA cycle is active, after feeding ^13^C_6_-glucose. Pink arrow: the reaction catalysed by succinate dehydrogenase (SDH), which converts succinate to fumarate. Isotopologues shown in green originate from the first cyclic TCA (M+2 acetyl-CoA and M+0 oxaloacetate). Isotopologues shown in blue are traditionally expected to be generated after the 4^th^ cycle, where preceding cycles led to a sequential build-up of labelled carbons in each TCA intermediate. (**B**) Illustration of isotopologues originating from the bifurcated TCA after the introduction of ^13^C_6_-glucose. M+3 pyruvate may be carboxylated using M+0 CO_2_, leading to M+3 oxaloacetate/aspartate (red), or M+1 CO_2_, leading to M+4 oxaloacetate/aspartate (orange). M+3 or M+4 oxaloacetate may condense with M+2 acetyl-CoA, leading to M+5 or M+6 citrate. The oxaloacetate and citrate isotopologues lead to illustrated succinate and glutamate in the bifurcated TCA cycle, as shown. A representation of decarboxylating reactions which can be the source of M+1 CO_2_ is included in this illustration. (**C**-**E**) Proportions of M+2, M+3, or M+4 fumarate relative to the entire pool of detected fumarate in *S. pombe* and *S. japonicus*. (**F**) Proportion of ^13^C-labelled glutamate within whole detected glutamate pool 30 min after ^13^C_6_-glucose addition. (**C-F**) Shown are means ±SEM of two biological and two technical replicates, *p* values were calculated using unpaired t-test. (**G**) Growth rates of *S. pombe* and *S. japonicus* grown in minimal EMM medium with 0.2g/L of either glutamate (E), glutamine (Q) or arginine (R). Means ±SEM of three technical and at least two biological replicates are shown, with p values generated using unpaired t-test.

M+2 fumarate is typically associated with the oxidative TCA cycle, as the two labelled carbons originate from acetyl-CoA condensing with unlabelled oxaloacetate (Fig. 2A, [50, 55-57]). Indeed, only respiro-fermenting wild type *S. pombe* showed M+2 fumarate (Fig. 2C). M+3 fumarate, on the other hand, likely originates from the reductive TCA branch, stemming from the carboxylation of M+3 pyruvate to oxaloacetate, which is then reductively converted to malate and fumarate (Fig. 2B, [50, 55-57]). Interestingly, both wild type and *cox6Δ S. pombe*, and wild type *S. japonicus* showed high M+3 fumarate fractions (Fig. 2D). This indicates that not only *S. japonicus* and non-respiring *S. pombe*, but also the wild type respiro-fermenting *S. pombe* operate the reductive TCA branch. However, the inability to respire in both fission yeasts leads to an increase in M+3 fumarate labelling, suggesting greater use of this bifurcated TCA architecture.

M+4 fumarate is often thought to result from repeated oxidative TCA cycles (Fig. 2A, [50, 55-57]). The wild type *S. pombe* indeed showed strong M+4 fumarate signal. However, this metabolite isotopologue was also highly abundant in both non-respiring *cox6Δ S. pombe* and the wild type *S. japonicus* (Fig. 2E). The proportion of M+4 fumarate in *S. pombe* was only minimally affected by the deletion of *sdh3*, which disrupts the cycle [58], suggesting that this signal largely did not originate from the oxidative TCA reactions. In fact, the relative abundances of M+4 fumarate in our data set were similar to the M+3 fractions, rather than the M+2 signal that was only present in the wild type *S. pombe* (Fig. 2C).

How is the abundant M+4 fumarate generated during fermentation? In principle, it could originate in reductive TCA reactions from the M+4 oxaloacetate (Fig. 2B). In turn, the M+4 oxaloacetate is likely synthesized through the carboxylation of the M+3 pyruvate by the pyruvate carboxylase using an M+1 labelled bicarbonate [59]. Supporting this hypothesis, we observed significant M+4 aspartate (proxy for oxaloacetate) labelling in both wild type fission yeasts, and their TCA cycle mutants (Fig. S2A). The labelled bicarbonate is likely released from decarboxylating reactions [59]. In fermenting cells one such reaction is the conversion of pyruvate to acetaldehyde [10]. Thus, a large fraction of M+4 fumarate even in respiro-fermenting cells could be a product of the reductive TCA branch and the recycling of endogenous bicarbonate.

Taken together, our genetic and metabolomics data suggest that 1) *S. japonicus* operates strictly a bifurcated TCA cycle with endogenous bicarbonate recycling; and 2) the respiro-fermenting *S. pombe* uses a combination of bifurcated and cyclic TCA reactions.

### Efficient NADH oxidation, but not cyclic TCA activity, is required for rapid glutamate synthesis

It has been postulated that efficient production of aKG-derived amino acids, such as glutamate and arginine, requires a canonical TCA cycle [18, 19, 60]. Seemingly in agreement with that hypothesis, glutamate production was slower in non-respiring *S. pombe cox6Δ* mutant as compared to the wild type (Fig. 2F). However, glutamate labelling was in fact more rapid in *S. japonicus* than in *S. pombe*, despite the lack of a canonical TCA cycle in the former (Fig. 2F). The *S. pombe sdh3Δ* mutant cells that have a broken TCA cycle synthesized glutamate similarly to the wild type (Fig. 2F). These results suggest that some aspect of respiration rather than running the canonical TCA cycle is important for efficient glutamate synthesis in *S. pombe*, whereas its sister species *S. japonicus* has evolved a respiration-independent strategy to boost amino acid production.

The oxidative branch of the TCA cycle generates NADH, which must be re-oxidised to support TCA reactions and biomass production. We wondered whether decreased NADH re-oxidation, rather than other defects in *S. pombe cox6Δ* mutant cells, was responsible for the reduced glutamate labelling. Ndi1 is the matrix-facing NADH dehydrogenase associated with the ETC. Deleting *ndi1* did not abolish respiration, as *S. pombe ndi1Δ* mutant cells grew on non-fermentable medium (Fig. S2B). Interestingly, *S. pombe ndi1Δ* cultures showed a pronounced reduction in glutamate labelling (Fig. 2F). Accordingly, supplementation with aKG-derived amino acids – glutamate, glutamine, or arginine – improved their growth in the minimal EMM medium (Fig. 2G). Surprisingly, the *S. pombe cox6Δ* mutants did not respond to glutamate or glutamine supplementation, and their growth defect in EMM could only be rescued by arginine (Fig. 2G). In contrast to what was suggested previously [18], this indicates that the arginine dependency of non-respiring *cox6Δ S. pombe* is not due to limited aKG production. *S. japonicus* appears to have overcome this limitation, as its growth is not improved by arginine supplementation (Fig. 2G).

Taken together, our results suggest that as long as cells can efficiently re-oxidize NADH generated by the oxidative TCA branch, a bifurcated TCA architecture can sustain efficient amino acid synthesis and biomass production.

### *S. japonicus* sustains higher glycolytic flux than *S. pombe*

Respiration is highly efficient for cellular ATP production, thanks to the additional oxidation of carbon substrates. We wondered whether *S. japonicus*’ inability to respire leads to a reduction in its overall energy content, or whether this species has optimized its glycolytic output (Fig. 3A). Surprisingly, *S. japonicus* exhibited a higher ATP/ADP ratio as compared to both respiro-fermenting (wild type) and solely fermenting (*cox6Δ*) *S. pombe* (Fig. 3B). This suggests that the energetic output of glycolysis in *S. japonicus* is higher than that of *S. pombe*.

**Figure 3.**
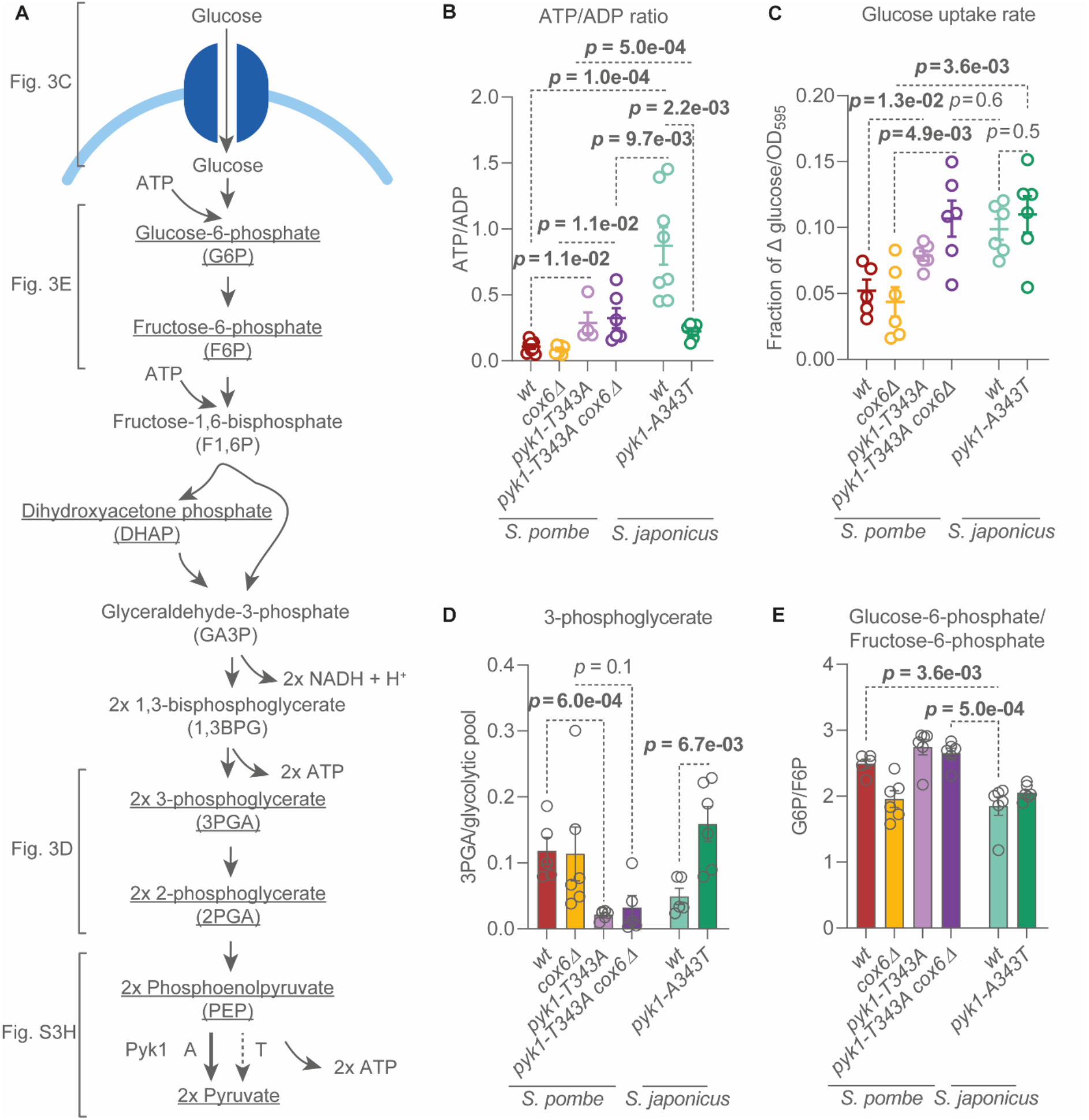
*S. japonicus* maintains higher glycolytic flux than *S. pombe*. (**A**) Illustration of glycolysis and its outputs for each molecule of glucose metabolized by the cell. Blue: a simplified example of a plasma membrane hexose transporter. Underlined are metabolites quantified in this study. Dephosphorylation of phosphoenolpyruvate by pyruvate kinase Pyk1 is indicated, with A and T denoting the point mutation at site 343 and its effects on Pyk1 activity. (**B**) ATP/ADP ratios detected in whole-cell extracts of *S. pombe* and *S. japonicus*. Means ±SEM values of at least four biological replicates. *p* values were calculated using unpaired t-test. (**C**) Rate of glucose uptake rate in EMM cultures of *S. pombe* and *S. japonicus*. Means ±SEM of at least five biological replicates are shown. *p* values were calculated using unpaired t-test. (**D**) 3-phosphoglycerate abundance relative to the sum of detected glycolytic intermediates (G6P, F6P, DHAP, 3PGA, 2PGA, PEP and pyruvate). (**E**) The abundance of glucose-6-phosphate relative to fructose-6-phosphate in whole-cell extracts of *S. pombe* and *S. japonicus*. (**D, E**) Plotted values are means ±SEM of three technical and two biological replicate experiments. *p* values were calculated using unpaired t-test.

Recent work has shown that the widely used laboratory ‘wild type’ strain of *S. pombe* possesses a point mutation in *pyk1*, the gene encoding pyruvate kinase that catalyses the last energy-yielding step of glycolysis (Fig. 3A). The alanine-to-threonine point mutation at position 343 (A343T) leads to a reduction in Pyk1 activity [61]. The Pyk1 kinase in *S. japonicus* does not have this partial loss-of-function substitution. To test if higher Pyk1 activity allows for more efficient glycolysis in this organism, we constructed a *pyk1-A343T S. japonicus* mutant and compared its ATP/ADP ratio with that of the wild type *S. japonicus*, wild type *S. pombe*, and *S. pombe* encoding a more active Pyk1 (*pyk1-T343A*). To hone specifically on the energetic output of glycolysis, we also constructed a non-respiring *S. pombe* strain with higher Pyk1 activity (*pyk1-T343A cox6Δ*).

Consistent with previous work [61], the higher Pyk1 activity in *S. pombe* increased the cellular ATP/ADP ratio, whereas the lower Pyk1 activity in *S. japonicus* reduced it (Fig. 3B). The ATP/ADP ratio of wild type *S. japonicus* was still higher than that of *pyk1-T343A cox6Δ S. pombe* (Fig. 3B). This result is notable because both cell types do not respire and presumably have the same level of pyruvate kinase activity.

In non-respiring cells, an improvement in ATP production could be due to an increase in glycolytic flux. We measured glucose uptake of exponentially growing cultures to estimate glycolytic rates in the two sister species [62, 63]. The higher activity of Pyk1 significantly improved glucose uptake in *S. pombe*, regardless of respiratory activity (Fig. 3C). The glucose uptake rate of *S. japonicus* was similar to that of the non-respiring *S. pombe* with high Pyk1 activity (*pyk1-T343A cox6Δ*). Interestingly, glucose uptake remained high in *S. japonicus* even when we introduced the *S. pombe*-specific partial loss-of-function *pyk1-A343T* allele (Fig. 3C). This suggests that unlike in *S. pombe*, the glycolytic flux in *S. japonicus* is not regulated as tightly by the pyruvate kinase. Notably, whereas the *pyk1* alleles do play a major role in modulating ATP production via glycolysis, *S. japonicus* might have evolved additional means of maximising glycolytic output.

To further probe the regulation of glycolysis in the two fission yeasts, we quantified glycolytic intermediates using GC-MS. The abundance of each intermediate was expressed as a fraction of the whole glycolytic intermediate pool, as this allowed us to pinpoint potential regulatory points (Fig. S3A-G). Corroborating published data, we observed a release in the PEP-to-pyruvate bottleneck when introducing a more active *pyk1* allele to *S. pombe* (Fig. S3H). Suggesting that the pyruvate kinase activity could indeed regulate glycolysis as a whole we have observed a Pyk1 activity-dependent bottleneck specifically at the 3PGA level in *S. pombe* (Fig. 3D). In line with the predicted higher pyruvate kinase activity in *S. japonicus*, we detected the accumulation of both PEP and 3PGA after introducing the *S. pombe*-like *pyk1-A343T* allele to this species (Fig. 3D and Fig. S3H).

Interestingly, regardless of the *pyk1* allele, *S. japonicus* exhibited a lower glucose-6-phosphate to fructose-6-phosphate (G6P/F6P) ratio than *S. pombe* (Fig. 3E). This suggested that G6P was consumed more rapidly in *S. japonicus*, highlighting potential differences in upper glycolysis between the two related organisms.

### *S. japonicus* may upregulate upper glycolysis via the pentose phosphate pathway

A major route utilizing upper glycolytic intermediates, and in particular, glucose-6-phosphate, is the pentose phosphate pathway (PPP) (Fig. 4A). To test whether the PPP was more active in *S. japonicus*, we have quantified the PPP intermediates in both fission yeasts. Consistent with rapid entry of G6P into the irreversible oxidative PPP, the glucose-6-phosphate to 6-phosphogluconate (G6P/6PGA) ratio was considerably lower in *S. japonicus* as compared to both wild type and solely fermenting *cox6Δ S. pombe* (Fig. 4B). This phenomenon was independent of the pyruvate kinase activity (Fig. 4B). Suggesting a higher flux through the oxidative PPP in *S. japonicus*, we detected a considerably higher NADPH / total NADP(H) as compared to *S. pombe* (Fig. 4C). As NADPH is a product of oxidative PPP (Fig. 4A), our results indicate that this pathway operates at higher capacity in *S. japonicus*. Suggesting a steady state conversion of 6PGA to Ru5P, the ratio of these metabolites in *S. japonicus* was close to 1 (Fig. S4A).

**Figure 4.**
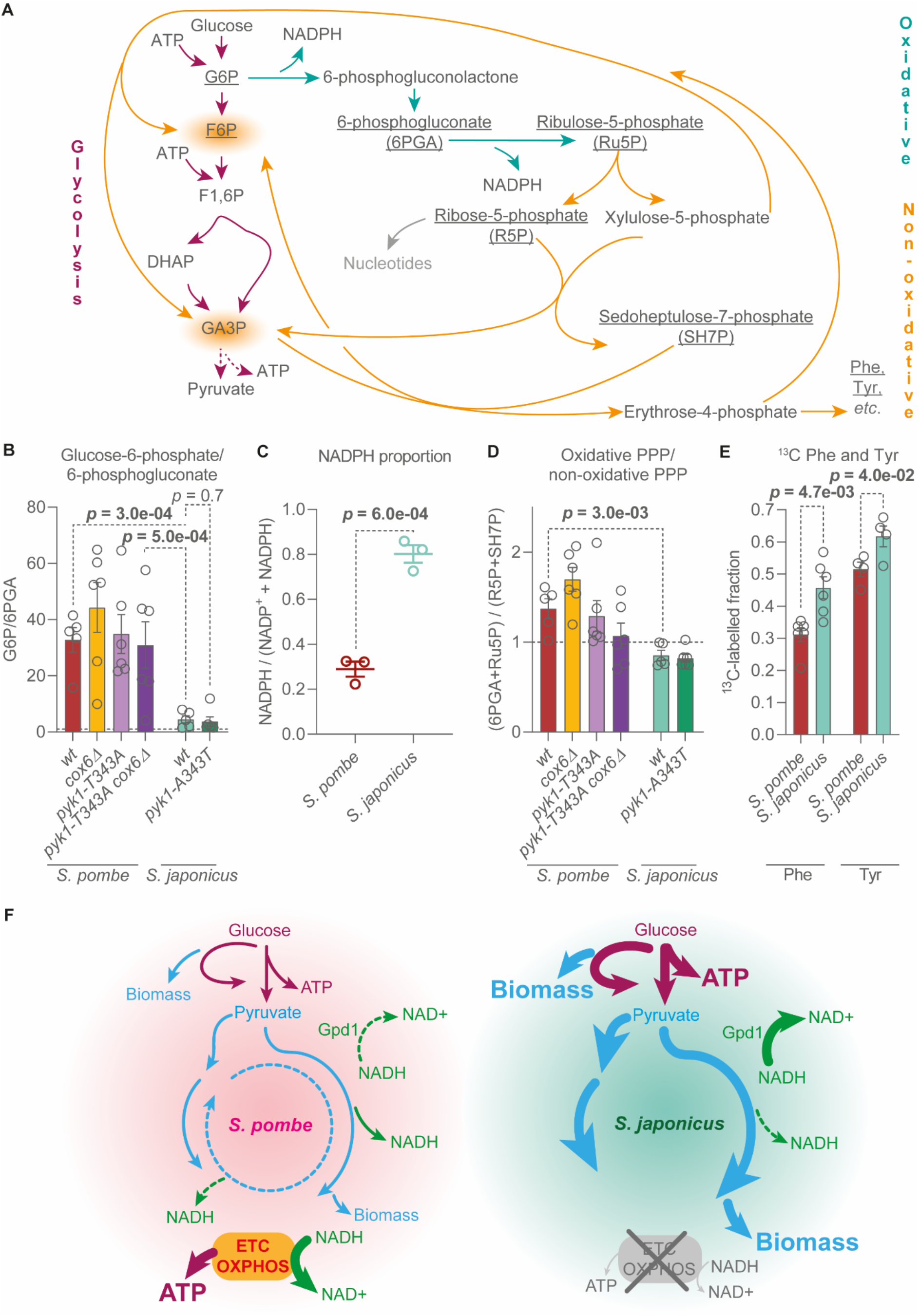
*S. japonicus* upregulates the pentose phosphate pathway as compared to *S. pombe*. (**A**) Illustration of the PPP and its intersection with glycolysis. Underlined are metabolites quantified in this study. G6P = glucose-6-phosphate, F6P = fructose-6-phosphate, F1,6BP = fructose-1,6-bisphosphate, DHAP = dihydroxyacetone phosphate, Phe = phenylalanine. Tyr = tyrosine. (**B**) Glucose-6-phosphate abundance normalised to 6-phosphogluconate. (**C**) Cellular NADPH relative to a total NADP(H) pool. Means ±SEM of three biological replicates, *p* values estimated using unpaired t-test. (**D**) Sum of oxidative PPP intermediates (6PGA and Ru5P) relative to non-oxidative PPP intermediates (R5P and SH7P). (**E**) Proportion of ^13^C-labelled phenylalanine and tyrosine within whole detected phenylalanine and tyrosine pools 10 min after ^13^C-labelled glucose addition. (**B, D**) Dotted lines indicate the ratio of 1. (**B, D, E**) Means ±SEM of two biological and two to three technical replicates. Statistical analyses were performed using unpaired t-test. (**F**) A diagram summarising the findings of this study on *S. pombe* and *S. japonicus* metabolism. ETC, electron transport chain, OXPHOS, oxidative phosphorylation.

The oxidative reactions of the PPP are followed by a non-oxidative part of the pathway. The latter allows cells to redirect carbon back into glycolysis (Fig. 4A, [64-66]). To understand if *S. japonicus* may use the PPP as a glycolytic shunt to maintain a higher glycolytic flux than *S. pombe*, we estimated the relative activities of oxidative vs non-oxidative PPP in both fission yeasts, by comparing the relative abundances of their intermediates. The ratio of oxidative relative to non-oxidative PPP products was lower in *S. japonicus*, consistent with increased non-oxidative PPP (Fig. 4D; see individual data for Ru5P/R5P and R5P/SH7P in Fig. S4B and S4C). Importantly, we observed faster biosynthesis of phenylalanine and tyrosine, amino acids produced from the non-oxidative PPP (Fig. 4A), in *S. japonicus* than *S. pombe* (Fig. 4E).

Taken together, these results suggest that a higher PPP activity in *S. japonicus* may allow this organism to maintain higher glycolytic flux as compared to *S. pombe*, in addition to producing more biomass and the essential anabolic cofactor NADPH.

## Discussion

Oxidative phosphorylation is the most efficient way of generating ATP per molecule of glucose. The ETC also oxidizes NADH to support anabolism. Indeed, the popular model fission yeast *S. pombe* relies on some degree of respiratory activity to grow at maximum capacity [13, 18]. Strikingly, its non-respiring relative *S. japonicus* grows rapidly, even when synthesizing all biomass precursors. This suggests that it has optimized its energy and redox metabolism, together with biomass production, in the absence of respiration (Fig. 4F).

First, *S. japonicus* appears to rely on cytosolic DHAP reduction to re-oxidize NADH (Fig. 1D, F). In addition to supporting amino acid and nucleotide anabolism, the high activity of glycerol-3-phosphate dehydrogenase Gpd1 presumably provides more glycerol backbones for lipid synthesis and results in higher yield of glycerol production. Interestingly, *S. japonicus* has lost the fructose-1,6-bisphosphatase Fbp1 [25, 67], rendering it incapable of performing gluconeogenesis, which may aid in supporting high glycolytic flux, and preventing glycerol consumption (Fig. S1A, B; see [68]). The loss of *fbp1* could allow *S. japonicus* to maintain abundant intracellular glycerol to potentially support high turgor pressure enabling rapid growth of this walled organism [34, 69].

Second, *S. japonicus* supports the production of amino acids derived from the TCA cycle by running a bifurcated version of this pathway (Fig. 2). Interestingly, even *S. pombe*, which requires respiration for optimal growth, runs both bifurcated and the canonical TCA cycle (Fig. 2). Presumably, the bifurcated version allows for better biomass production by committing oxaloacetate and aKG to amino acid synthesis. Our data on the *S. pombe ndi1* mutant demonstrates that the bifurcated version of the TCA cycle yielding biomass still depends on efficient ETC-dependent NADH oxidation in this species (Fig. 2F, G). By utilizing the bifurcated TCA alongside respiration, *S. pombe* maintains rapid biomass production and growth. Yet, the fast growth of *S. japonicus* demonstrates that this is not the only effective strategy. In addition to optimizing ETC-independent NADH oxidation, *S. japonicus* appears to have rewired the requirements for the synthesis of aKG-derived amino acids. It has lost both subunits of the NAD-dependent isocitrate dehydrogenase IDH, while retaining the NADP-dependent isocitrate dehydrogenase Idp1. In addition, it has lost one of the NAD-dependent glutamate dehydrogenases Gdh2 but kept the NADP-specific glutamate dehydrogenase Gdh1 [25, 67, 70, 71]. This suggests that besides evolving efficient NADH oxidation pathway(s), *S. japonicus* has alleviated the NADH burden by changing the cofactor dependencies of anabolic enzymes.

Third, *S. japonicus* maintains high ATP levels by maximizing glycolytic flux through a combination of high pyruvate kinase activity and bypassing an initial energy consuming step of glycolysis via the PPP (Fig. 4). Diverting glucose-6-phosphate away from glycolysis and reintroducing carbon back at the glyceraldehyde-3-phosphate level may reduce allosteric inhibition of glycolytic enzymes such as hexokinase [72], major glycolytic checkpoint enzymes such as phosphofructokinase [63, 73], and avoid one of the ATP-demanding steps of upper glycolysis. This behaviour appears to be upregulated in *S. japonicus*, as compared to *S. pombe*.

Although the replacement of *S. japonicus pyk1* with the *S. pombe*-like allele encoding a pyruvate kinase with lower activity led to the accumulation of phosphoenolpyruvate (Fig. S3H) and 3-phosphoglycerate (Fig. 3D), *S. japonicus* glucose uptake was not reduced (Fig. 3C). This suggests that *S. japonicus* upper glycolysis is less tightly regulated by the flux through lower glycolysis, as compared to its sister species. Presumably, this feature allows *S. japonicus* to sustain rapid glycolysis in situations when it normally would be inhibited, such as low pH [74, 75], or low environmental glucose levels [76-78]. The latter could be particularly important, since *S. japonicus* cannot rely on respiration as a metabolic strategy for dealing with starvation, unlike *S. pombe* [12, 13, 54]. The lack of Fbp1 may also prevent the suppression of glycolysis in low glucose [79], which is arguably vital for a non-respiring organism. Importantly, the lack of gluconeogenesis in *S. japonicus* may support the use of the non-oxidative PPP as a one-way shunt into glycolysis, and would prevent the recycling of F6P and GA3P into the oxidative PPP [65]. Overall, a combination of upregulated PPP, ‘good’ pyruvate kinase, and other adaptations reducing negative feedback on glycolysis, allows *S. japonicus* to maintain high ATP levels independent of OXPHOS.

Finally, the upregulation of the pentose phosphate pathway may also allow *S. japonicus* to rapidly produce nucleotides and some amino acids, such as phenylalanine and tyrosine (Fig. 4E). Furthermore, it results in high yields of NADPH (Fig. 4C), which is key for several anabolic pathways, e.g., lipid metabolism [80]. *S. japonicus* is highly sensitive to paraquat and hydrogen peroxide [26], despite high levels of NADPH, which is needed for the reduction of glutathione and thioredoxin [81, 82]. It is plausible that over the course of its life history, which has involved adaptation to anaerobic environment, this species has lost some capacity to manage oxidative stress.

Our study lays the groundwork for better understanding of central carbon metabolism in fission yeasts and beyond. We demonstrate the power of stable isotope tracing metabolomics, mass isotopologue distribution analyses and genetic perturbations in illuminating the architecture of metabolic pathways in yeasts. Importantly, our work showcases a comparative biology approach to understanding metabolism.

Central carbon metabolism is woven tightly into the fabric of cellular biology. Understanding the plasticity of metabolism both in ontogenetic and phylogenetic terms may ultimately aid in explaining organismal ecology and the evolution of higher-level cellular features, such cell size and growth rate.

## Materials and methods

### Maintenance and growth of fission yeasts

*S. pombe* and *S. japonicus* prototrophic strains used in this study are listed in Table S1. Standard fission yeast media and methods were used [83, 84]. For non-fermentable conditions, we used EMM with 2% glycerol or 2% galactose and 0.1% glucose. The inclusion of 0.1% glucose is necessary for *S. pombe* growth in these conditions [54, 85]. For most experiments, yeasts were pre-cultured in Edinburgh Minimal Medium (EMM) at 30°C. Pre-cultures for growth, serial dilution or oxygen consumption experiments were grown in rich Yeast Extract and Supplements (YES) medium at 24°C. All cultures were grown in a 200rpm shaking incubator. The following day, cultures were diluted to an OD_595_ within lag-phase or early-exponential phase, as required, and allowed to grow to a desired OD_595_. Once cultures reached early (0.2-0.5 OD_595_) or mid-exponential phase (OD_595_ determined by each strain and condition’s OD_595_ at stationary phase), cells were collected.

In the case of growth experiments, post-dilution growth was tracked in a plate reader, as follows. Yeasts were pre-cultured in YES at 25°C until OD_595_ 0.1-0.6. Cultures were then washed in experimental medium and diluted to 0.1 OD_595_. Growth was measured every 10 minutes at 30°C using VICTOR Nivo multimode plate reader (PerkinElmer). Growth curves were plotted using Graphpad Prism and growth rates were calculated using the Growthcurver R package [86]. All experiments were performed in three technical and at least three biological replicates. Technical replicates were three wells in a 96-well plate; biological replicates were independent growth experiments using freshly-defrosted batches of each strain.

### Serial dilution assays

Serial dilution assays were performed by preculturing cells in YES at 25°C overnight until early-xponential phase. Cultures were diluted to 2×10^6^ cells/ml and serially diluted by a factor of 10. 2µl of each dilution were inoculated on plates. Plates were typically incubated at 30°C for three days. Anaerocult P (Millipore) anoxic bags were used for anaerobic growth. All experiments were repeated on three separate occasions, using freshly-defrosted strains.

### Hyphae formation assay

*S. japonicus* cultures grown at 24°C overnight in YES to exponential phase were centrifuged and washed once in YES. The equivalent of OD_595_ 0.5 was inoculated on Yeast Extract Glucose Malt Extract Agar (YEMA) plates and incubated at 30°C for five days. Surface colonies were gently washed away to retain only the hyphae formed within the agar. Plates were imaged and Image J was used to measure the diameter of hyphae formed. Experiments were repeated at least thrice using freshly defrosted strains.

### Sporulation efficiency assay

*S. japonicus* wild-type and deletion strains of the opposite mating type, carrying the kanMX (KanR) selection marker, were crossed on Sporulation Agar (SPA) plates overnight at 30°C. The following day, a defined numbers of spores were dissected on YES plates using a Singer MSM micromanipulator (Singer Instruments). Between 50 and 100 spores per biological replicate were dissected. Spores were allowed to germinate over three days at 30°C. Spores that formed colonies were counted to estimate sporulation efficiency. Colonies of spores from wild-type and deletion strain crosses were replica-plated onto G418 sulphate (Sigma Aldrich) containing YES plates to determine the KanMX positivity score. Experiments were replicated at least thrice independently.

### Strains and molecular biology methods

Molecular genetic manipulations were performed via homologous recombination using the gene deletion cassette method [87, 88], where target genes’ open reading frames were replaced by a pFA6A-based plasmid with kanR, NatR or HygR cassettes flanked by 80-base-pair portions of the 5’ and 3’ UTRs of the gene of interest.

*S. pombe* was transformed using the lithium acetate method, as previously described [89]. Briefly, early-exponential *S. pombe* cultures grown in YES were centrifuged and washed twice in dH_2_O. Cells were then washed in lithium acetate Tris-EDTA and incubated in 100µl of the same buffer for 10 minutes at room temperature together with 5µg of linear DNA and 50µg of sonicated salmon sperm DNA (Agilent Technologies). 240µl of PEG-lithium acetate Tris-EDTA was added and cell suspension was mixed by swirling with a pipette tip. Samples were incubated at 30°C for 30-60 minutes. 43µl of DMSO was then added and cells were washed in dH_2_O twice. Cells were then recovered in 10ml of YES at 25°C overnight and subsequently inoculated on selective plates - YES plates containing 100µg/mL G418 sulphate (Sigma Aldrich), 50µg/mL hygromycin B (Sigma Aldrich), or 100µg/mL nourseothricin (Werner BioAgents, Germany).

*S. japonicus* was transformed using electroporation [84]. Briefly, early-exponential cultures grown in YES were pelleted and from then on kept on ice. Cells were washed three times using ice-cold dH_2_O and then were suspended in 5ml of cold 1M sorbitol with 50mM dithiothreitol (DTT) in dH_2_O. Suspensions were incubated for 12 minutes at 30°C (without shaking) after which cells were centrifuged and washed twice with cold 1M sorbitol. Pellets were then resuspended in 100µl 1M sorbitol containing linearised DNA and sonicated salmon sperm DNA (Agilent Technologies). Cells were incubated on ice for 30 minutes. *S. japonicus* was subsequently electroporated at 2.30keV using a cold 2mm Gene Pulser/MicroPulser Electroporation Cuvette (Bio-Rad laboratories). Immediately after electroporation, 1ml of cold 1M sorbitol was added to the cell suspension, and cells were recovered overnight at 25°C in 10ml YES, shaking. The next day, cells were plated on selective plates, as with *S. pombe*.

### Oxygen consumption

Oxygen levels were measured using a waterproof field Dissolved Oxygen meter (Hanna HI 98193). Oxygen consumption was measured in mid-exponential cultures grown in YES. Cultures were centrifuged and resuspended in fresh medium and used to fill a conical flask to the brim. The probe was submerged into the culture and the flask was sealed. Cultures were kept in gentle motion using a magnetic stirrer. Once oxygen readings stabilised oxygen levels were recorded every minute for 10 minutes, after which the OD_595_ of cultures were recorded. Cultures were at room temperature during oxygen readings. Oxygen concentration over the time course was plotted to identify when oxygen changes slowed or plateaued. Selected linear oxygen changes were used to calculate the oxygen consumption rate per minute, which were normalised to the culture OD_595_. Measurements were independently repeated three times to yield three biological replicates.

### Glucose consumption

Cells were pre-cultured in EMM with 2% glucose and once cultures reached early exponential phase, they were diluted to OD_595_ 0.1 and incubated overnight at 30°C. When cultures reached mid-exponential phase, cells were centrifuged and resuspended in fresh medium. Cultures were placed in a shaking incubator at 30°C and media samples were collected after two hours. The decrease in glucose was normalised to the change in the OD_595_. Glucose was quantified using Glucose (HK) Assay Kit (Sigma Aldrich) as per the manufacturer’s instructions. Samples were collected as independent biological replicates from cultures grown on separate occasions, using freshly-defrosted strains.

### NAD(P)+/NAD(P)H quantification

Cells were pre-cultured in EMM overnight at 30°C until exponential phase. Subsequently, cultures were diluted to lag-phase or early-exponential phase and were grown overnight at 30°C. The equivalent of OD_595_5 of early-exponential (in the case of NAD^+^/NADH) or mid-exponential (in the case of NADP^+^/NADPH) cultures were harvested by centrifugation and snap-freezing in liquid nitrogen. NAD^+^/NADH was extracted and measured using MAK037 (Sigma Aldrich) and NADP^+^/NADPH was extracted and measured using MAK038 (Sigma Aldrich) as per manufacturer’s instructions. Cells were resuspended in chilled extraction buffer and lysed using lysing matrix Y tubes (MP Biomedicals) containing 0.5 mm diameter yttria-stabilized zirconium oxide beads and a cell disruptor. Samples were kept as cold as possible by bead beating in 10 second intervals, at 6.5m/sec, 10 times, with 2-minute breaks on ice between each round. Samples were then filtered through a 10kDa protein filter (MRCPRT010, Sigma Aldrich) via centrifugation at 4°C. Extracts were then processed as per the manufacturer’s instructions. Samples were collected in at least three independent experiments. Quantification was performed using a Tecan Spark plate reader.

### ATP/ADP quantification

Early-exponential cultures in EMM were collected by quenching the equivalent of OD_595_5 cells in -80°C methanol. Suspensions were centrifuged at 3000rpm for 2 minutes at 4°C and decanted, and pellets were dried in -80°C overnight. ATP and ADP were extracted and quantified using the ATP/ADP ratio quantification kit (MAK-135, Sigma Aldrich) as per the manufacturer’s instructions, with the following modification. To extract ATP and ADP, cell pellets were lysed in the kit’s assay buffer using lysing matrix Y tubes (MP Biomedicals) containing 0.5 mm diameter yttria-stabilized zirconium oxide beads and a cell disruptor. Samples were kept as cold as possible by bead beating once for 10 seconds at 6.5m/sec at 4°C. Bioluminescence was quantified using a Tecan Spark microplate reader. Samples were collected as at least three biological replicates from cultures grown on separate occasions, using freshly-defrosted strains.

### Gas Chromatography-Mass Spectrometry Metabolomics

Unprocessed integrals derived from all metabolomics experiments are shown in Tables S2-S6. The abundances of targeted metabolites in standard metabolite mix used in all experiments are listed in Table S7.

For metabolomics experiments, cells were pre-cultured in EMM and diluted the previous day so that cells were in early-exponential phase at the time of harvest. In the case of stable isotope tracing experiments, pre-cultures and experimental cultures were both grown in EMM at 24°C. When cultures reached an OD_595_ of 0.2-0.4, cells were centrifuged at 3000rpm for two minutes and resuspended in either unlabelled (^12^C) or labelled (^13^C) media without dilution. Labelled media refers to EMM with 2% D-Glucose (U-^13^C_6_) (Cambridge Isotope Laboratories, Inc.). Once cells came into contact with labelled media, a time course was started. Cells were kept agitated until each time point was reached. At 1, 3, 5, 10, 30 or 60 minutes, a volume equivalent to 1.5OD_595_ was injected into 100% LCMS-grade methanol (Sigma Aldrich) pre-cooled to -80°C. Quenched cultures were centrifuged and washed twice with -80°C methanol, centrifuging at 4°C and 3000rpm. The dried pellets stored at -80°C until extraction. For abundance quantification, a total of six replicates were collected per condition, two technical over three biological repeats. For stable isotope tracing, a total of four replicates were collected, two biological repeats with two technical replicates. Biological replicates were defined as independent experiments performed using freshly-defrosted batches of cells.

The extraction protocol is a modified version of a method developed in [19, 90]. Cell pellets were resuspended in 200µl of LCMS-grade acetonitrile/methanol/water (2:2:1) (Sigma Aldrich) chilled to -20°C and transferred to lysing matrix Y tubes (MP Biomedicals) containing 0.5 mm diameter yttria-stabilized zirconium oxide beads. Extraction blanks were included from this stage, consisting of 200µl of extraction solution. 1nmol of scyllo-inositol (Sigma Aldrich) standard was added to all samples at this stage. Samples were kept on ice and lysed at 4°C. Bead beating was performed at 6.5m/sec for 10 seconds, five times, with two-minute breaks on ice after each round. Subsequently, samples were centrifuged at 12,000rpm for two minutes at 4°C and supernatants were dried for 1-2 hours in a SpeedVac Vacuum Concentrator at 30°C. Samples were then stored at -80°C.

Dried extracts were resuspended in -20°C chilled 50µl LCMS-grade chloroform (Sigma Aldrich) and 300µl LCMS-grade methanol:water (1:1) (Sigma Aldrich). Samples were vortexed for one minute and centrifuged at 12,000rpm for five minutes at 4°C. 240µl of upper, polar phase was transferred to GC-MS glass vial inserts for drying, which included two 30µl methanol washes to ensure there was no residual water.

Derivatisation was performed based on published work [49]. Samples were resuspended in 20µl of 20mg/ml of freshly dissolved methoxyamine hydrochloride (Sigma Aldrich) in pyridine (Sigma Aldrich). Samples were briefly vortexed and centrifuged, and incubated at room temperature overnight (around 15 hours). The next day, 20µl of room-temperature N,O-bis(trimetylsilyl)trifluoroacetamide (BSTFA) and 1% trimethylchlorosilane (TMCS) (Sigma Aldrich) was added to each sample, followed by a brief vortex and centrifugation.

Metabolites were detected using Agilent 7890B-MS7000C GC-MS as previously described [49]. Prior to use, an MS tune was performed. Samples were arranged and processed in random order. Each batch contained six metabolite mixes (which functioned as quality control in addition to aiding metabolite identification and quantification) arranged equally at the start, middle and end. Hexane washes were run at the beginning and end of each run, between every six samples and before and after each metabolite mix. Splitless injection was performed at 270°C in a 30m + 10m x 0.25mm DB-5MS+DG column (Agilent J&W). Helium was used as the carrier gas. The oven temperature cycle was as follows: 70°C for 2 minutes, then a temperature gradient was applied with temperatures incrementing at a rate of 12.5°C/minute, reaching a maximum of 295°C, followed by a gradient of 25°C/minute reaching a maximum of 350°C. The 350°C temperature was held for 3 minutes. Electron impact ionization mode was used for MS analysis.

Samples were analysed using a combination of MassHunter Workstation (Agilent Technologies) and MANIC, an updated version of the software GAVIN [91], for identification and integration of defined ion fragment peaks. For abundance quantification, integrals, the known amount of scyllo-inositol internal standard (1nmol) and the known abundances in the standardised metabolite mix run in parallel to samples (Table S7) were used to calculate an estimated nmol abundance of each metabolite in each sample. The formula used for calculating molar abundances as shown in Equation 1. Abundances were plotted as relative proportions of a pool of related metabolites to facilitate cross-species and cross-condition comparisons.

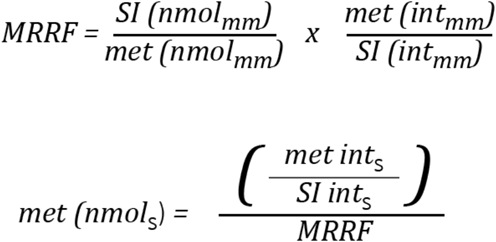

**Equation 1 - Formula for nmol abundance quantification**.

MRRF = molar relative response factor; SI = scyllo-inositol (internal standard); met = metabolite to be quantified; mm = standard metabolite mix; int = integrals; s = samples.

### Statistical analyses

All plots were generated using Graphpad Prism. All data was analysed using unpaired t-test statistical analysis, unless indicated otherwise.

## Acknowledgements

We are grateful to the Oliferenko and Bähler labs for discussions and Eugene Makeyev, Alice Yuen and Eneko Pascual Navarro for suggestions on the manuscript. We thank Anna Forbes for assistance with generating the *pyk1 S. japonicus* mutant strain. We are grateful to Ken Ishikawa (National Cancer Institute, U.S.) for the *S. japonicus var versatilis* strain, Matthew L. Bochman (Indiana University Bloomington, U.S.) and Makoto Kawamukai (Shimane University, Japan) for *S. japonicus* wild isolate strains. Many thanks to James MacRae and James Ellis (Francis Crick Institute Metabolomics Science Technology Platform) for invaluable training and assistance with metabolomics experiments. This research was funded in whole, or in part, by the Wellcome Trust (103741/Z/14/Z; 220790/Z/20/Z). For the purpose of Open Access, the author has applied a CC-BY public copyright licence to any Author Accepted Manuscript version arising from this submission.

## Author contributions

Sara Alam conceived, performed and interpreted metabolomics, physiology, and most genetics experiments; generated strains; analysed data; and co-wrote the manuscript. Ying Gu generated *gpd1Δ* and *mdh1Δ S. pombe* and *S. japonicus* strains, designed the *pyk1-A343T S. japonicus* mutant, and edited the manuscript. Polina Reichert assisted with analysis and interpretation of metabolomics experiments and edited the manuscript. Jürg Bähler interpreted experiments and edited the manuscript. Snezhana Oliferenko conceived and interpreted experiments, co-wrote and edited the manuscript.

## Funding

Sara Alam was supported by the Crick-King’s PhD scholarship. Work in S.O. lab was supported by the Francis Crick Institute, which receives its core funding from Cancer Research UK (CC0102), the UK Medical Research Council (CC0102), and the Wellcome Trust (CC0102), and the Wellcome Trust Senior Investigator Award (103741/Z/14/Z), Wellcome Trust Investigator Award in Science (220790/Z/20/Z) and BBSRC (BB/T000481/1) to Snezhana Oliferenko.

## Competing interests

The authors declare no competing or financial interests.

**Table S1.**
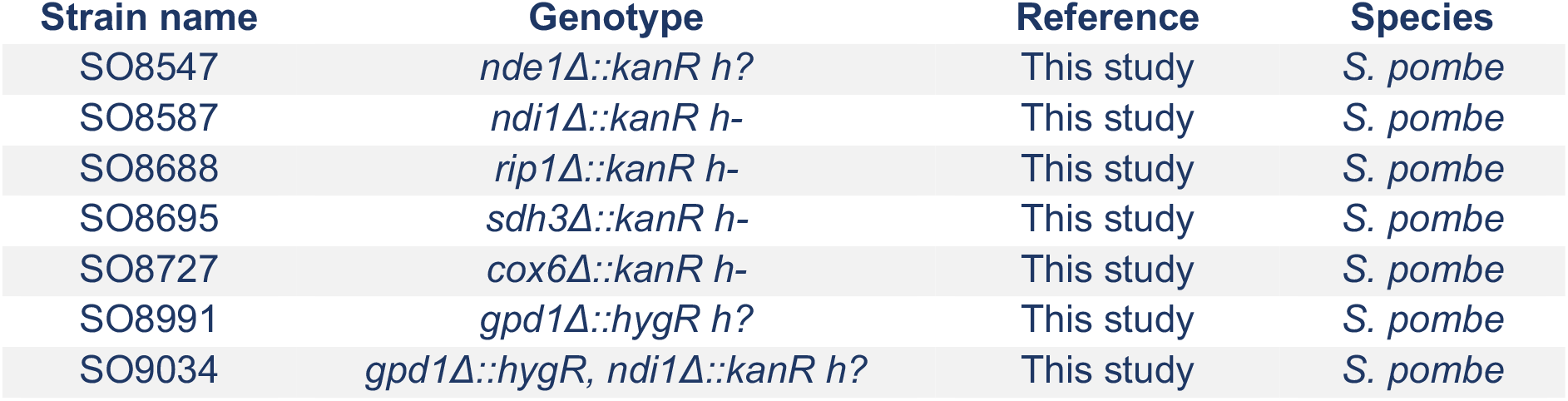
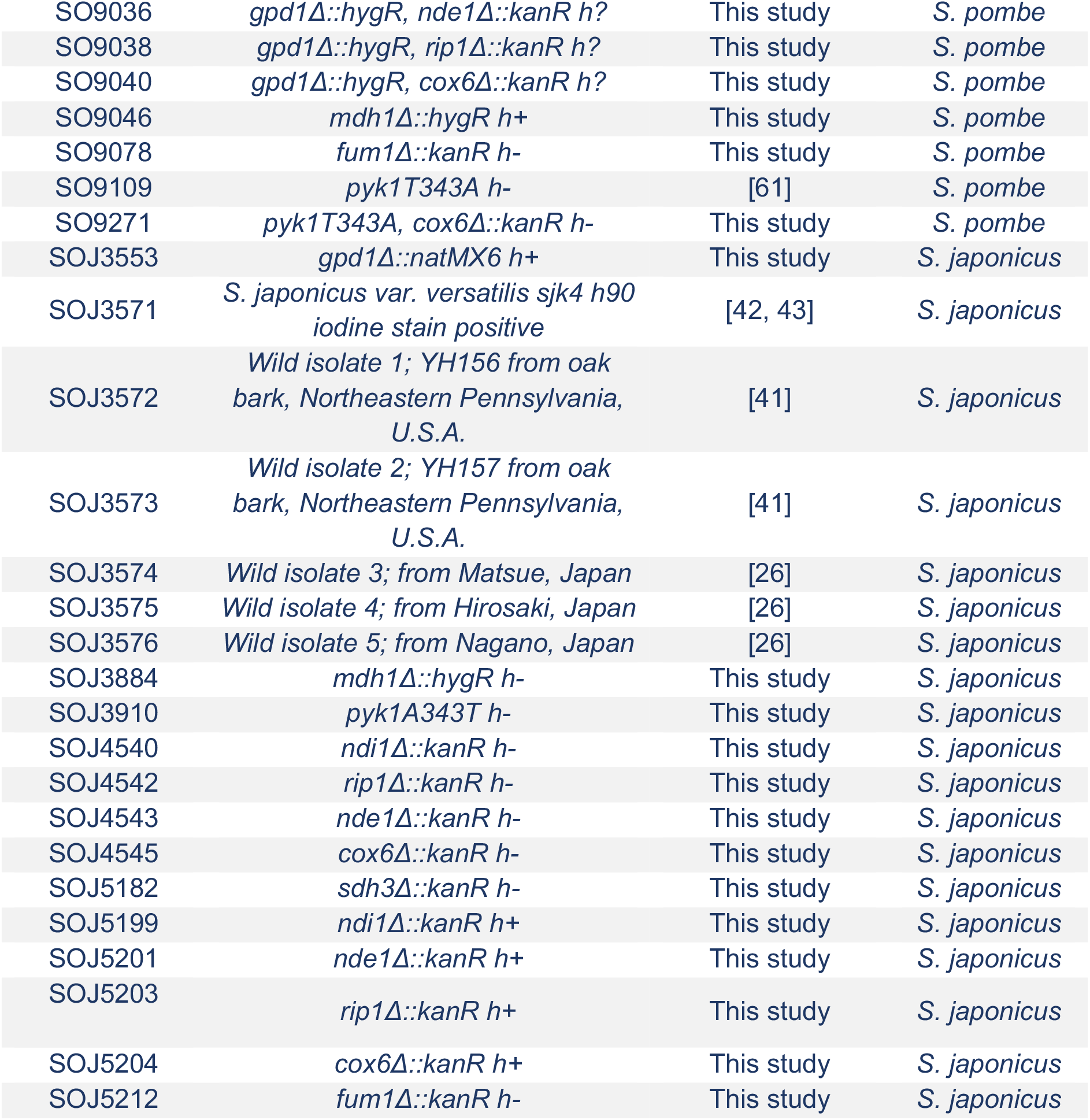
*S. pombe* and *S. japonicus* strains used in this study.

| Strain name | Genotype | Reference | Species |
| --- | --- | --- | --- |
| SO8547 | *nde1Δ::kanR h?* | This study | *S. pombe* |
| SO8587 | *ndi1Δ::kanR h-* | This study | *S. pombe* |
| SO8688 | *rip1Δ::kanR h-* | This study | *S. pombe* |
| SO8695 | *sdh3Δ::kanR h-* | This study | *S. pombe* |
| SO8727 | *cox6Δ::kanR h-* | This study | *S. pombe* |
| SO8991 | *gpd1Δ::hygR h?* | This study | *S. pombe* |
| SO9034 | *gpd1Δ::hygR, ndi1Δ::kanR h?* | This study | *S. pombe* |
| SO9036 | *gpd1Δ::hygR, nde1Δ::kanR h?* | This study | *S. pombe* |
| SO9038 | *gpd1Δ::hygR, rip1Δ::kanR h?* | This study | *S. pombe* |
| SO9040 | *gpd1Δ::hygR, cox6Δ::kanR h?* | This study | *S. pombe* |
| SO9046 | *mdh1Δ::hygR h+* | This study | *S. pombe* |
| SO9078 | *fum1Δ::kanR h-* | This study | *S. pombe* |
| SO9109 | *pyk1T343A h-* | [61] | *S. pombe* |
| SO9271 | *pyk1T343A, cox6Δ::kanR h-* | This study | *S. pombe* |
| SOJ3553 | *gpd1Δ::natMX6 h+* | This study | *S. japonicus* |
| SOJ3571 | *S. japonicus* var. *versatilis* *sjk4 h90*<br>iodine stain positive | [42, 43] | *S. japonicus* |
| SOJ3572 | Wild isolate 1; YH156 from oak<br>bark, Northeastern Pennsylvania,<br>U.S.A. | [41] | *S. japonicus* |
| SOJ3573 | Wild isolate 2; YH157 from oak<br>bark, Northeastern Pennsylvania,<br>U.S.A. | [41] | *S. japonicus* |
| SOJ3574 | Wild isolate 3; from Matsue, Japan | [26] | *S. japonicus* |
| SOJ3575 | Wild isolate 4; from Hirosaki, Japan | [26] | *S. japonicus* |
| SOJ3576 | Wild isolate 5; from Nagano, Japan | [26] | *S. japonicus* |
| SOJ3884 | *mdh1Δ::hygR h-* | This study | *S. japonicus* |
| SOJ3910 | *pyk1A343T h-* | This study | *S. japonicus* |
| SOJ4540 | *ndi1Δ::kanR h-* | This study | *S. japonicus* |
| SOJ4542 | *rip1Δ::kanR h-* | This study | *S. japonicus* |
| SOJ4543 | *nde1Δ::kanR h-* | This study | *S. japonicus* |
| SOJ4545 | *cox6Δ::kanR h-* | This study | *S. japonicus* |
| SOJ5182 | *sdh3Δ::kanR h-* | This study | *S. japonicus* |
| SOJ5199 | *ndi1Δ::kanR h+* | This study | *S. japonicus* |
| SOJ5201 | *nde1Δ::kanR h+* | This study | *S. japonicus* |
| SOJ5203 | *rip1Δ::kanR h+* | This study | *S. japonicus* |
| SOJ5204 | *cox6Δ::kanR h+* | This study | *S. japonicus* |
| SOJ5212 | *fum1Δ::kanR h-* | This study | *S. japonicus* |

---

**Supplemental Figure 1, related to Figure 1.**
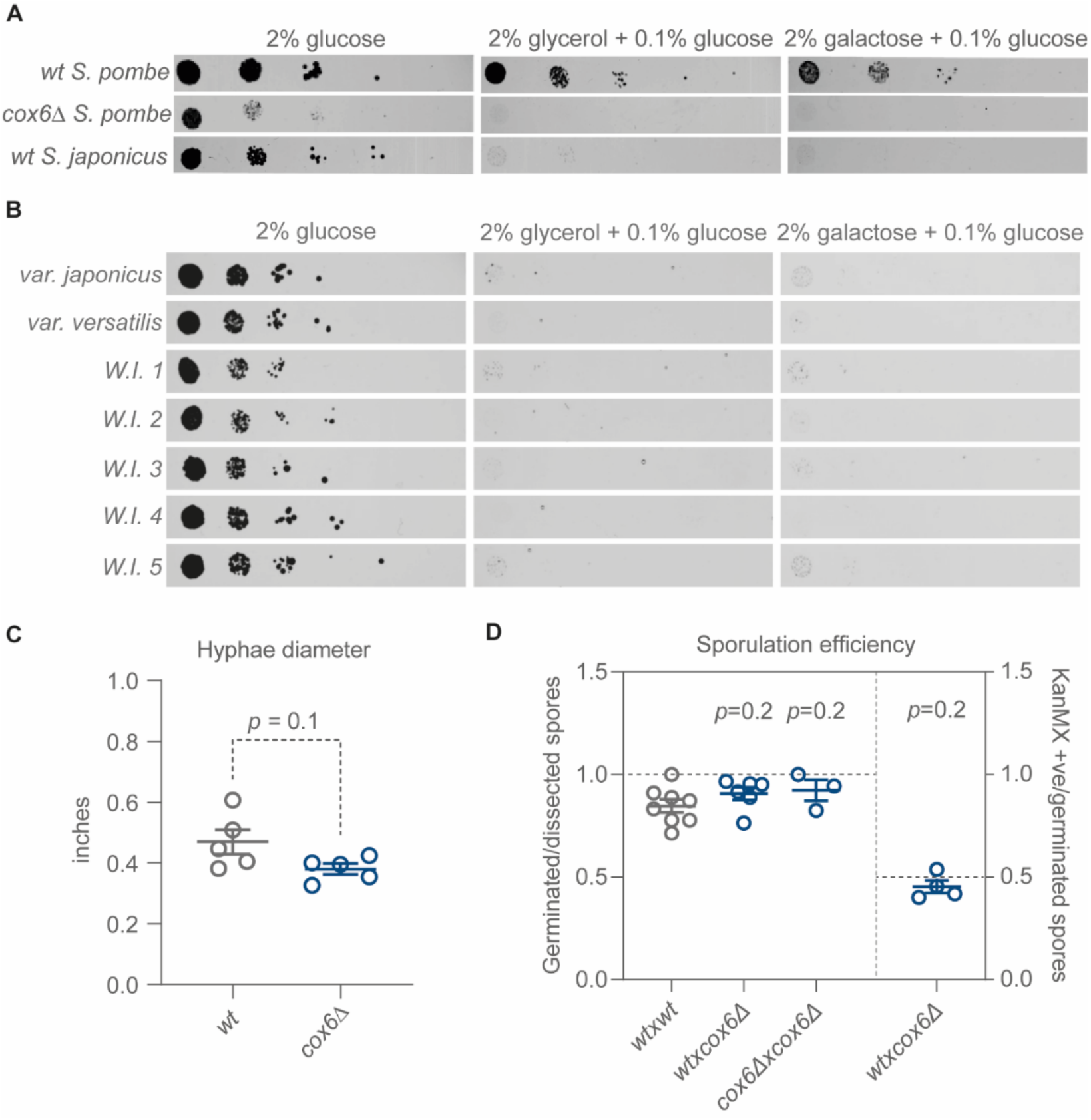
(**A**) Serial dilution assays showing growth of *S. pombe* and *S. japonicus* of indicated genotypes on EMM agar with either 2% w/v glucose, 2% v/v glycerol or 2% w/v galactose with 0.1% w/v glucose. Serially diluted strains were grown over four days at 30°C, images are representative of three independent biological replicates. (**B**) Serial dilution assays showing growth of wild type *S. japonicus* strains on EMM, as in (**A**). *var japonicus* is our wild type lab strain. *var versatilis* is a lab strain from the USA. W.I, wild isolate; W.I.1: YH156 from oak bark, Northeastern Pennsylvania, U.S.A.; W.I.2: YH157 from oak bark, Northeastern Pennsylvania, U.S.A.; W.I.3-5: isolates from Matsue, Hirosaki and Nagano, Japan. (**C**) Hyphae diameters measured five days after inoculation on YEMA (yeast extract, glucose and malt extract agar) plates. Plots represent means ±SEM of two biological and two to three technical replicates. *p* values were generated using unpaired t-test. (**D**) *Left*, sporulation efficiency of *S. japonicus* crosses of indicated genotypes, calculated by counting germinated spores proportionally to total dissected spores. *Right*, the recovery of *cox6Δ* KanMX-marked progeny did not deviate from the expected 50%. Spore germination score *p* values were generated using unpaired t-test, KanMX positivity score *p* values were calculated using one sample t-test assuming a Gaussian distribution, testing against a hypothetical mean of 0.5. Plots represent means ±SEM of two technical and two or three biological replicates.

**Supplemental Figure 2, related to Figure 2.**
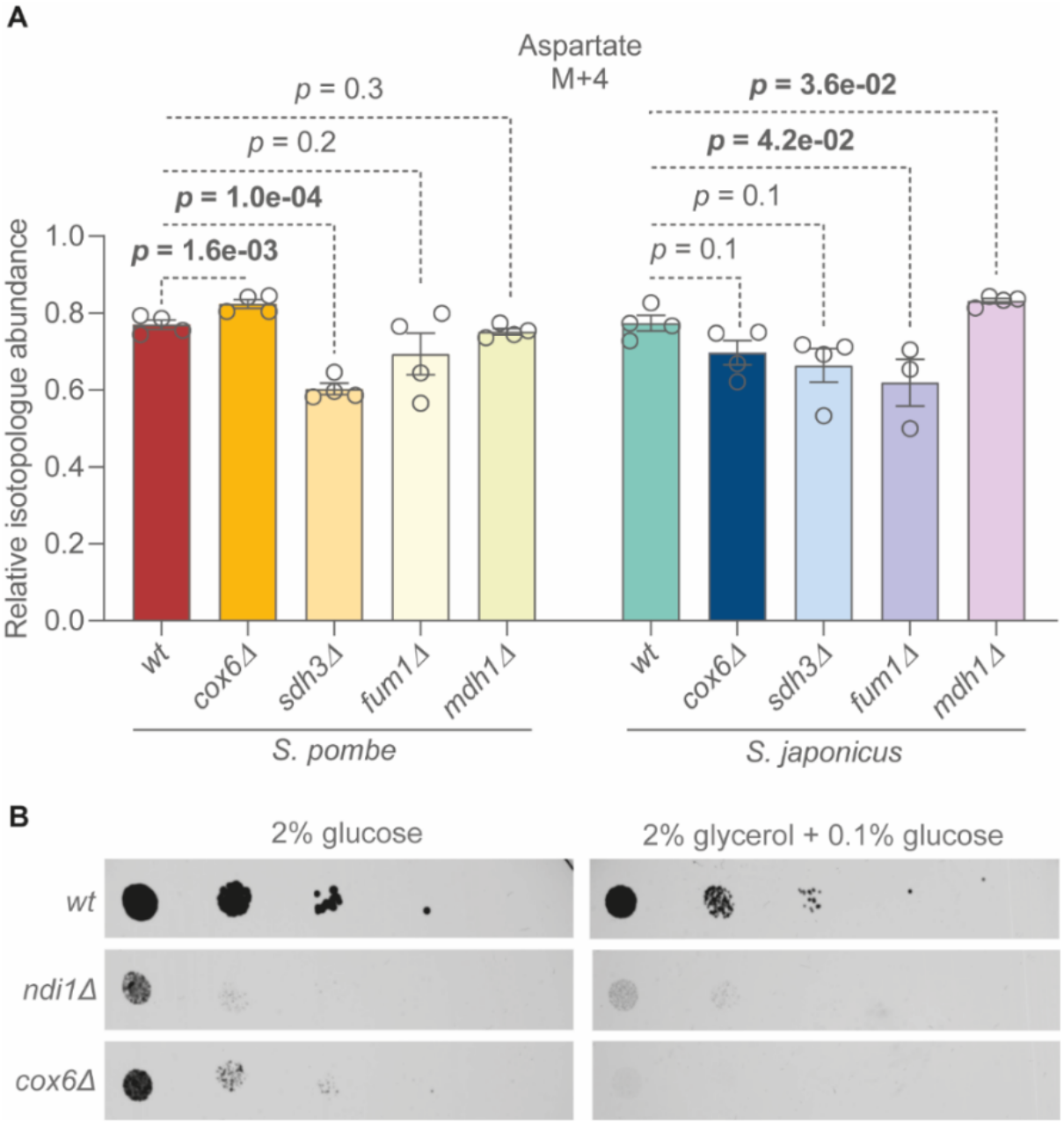
(**A**) Proportion of M+4 aspartate (proxy for oxaloacetate) relative to the detected pool of aspartate in *S. pombe* and *S. japonicus* 30 minutes after ^13^C_6_ glucose addition. Means ±SEM of two biological and two technical replicates, p values calculated using unpaired t-test. (**B**) Serial dilution assays of *S. pombe* on EMM agar plates with either 2% (w/v) glucose or 2% (v/v) glycerol with 0.1% (w/v) glucose. Serially diluted strains were grown over four days at 30°C, images are representative of three independent biological replicates.

**Supplemental Figure 3, related to Figure 3.**
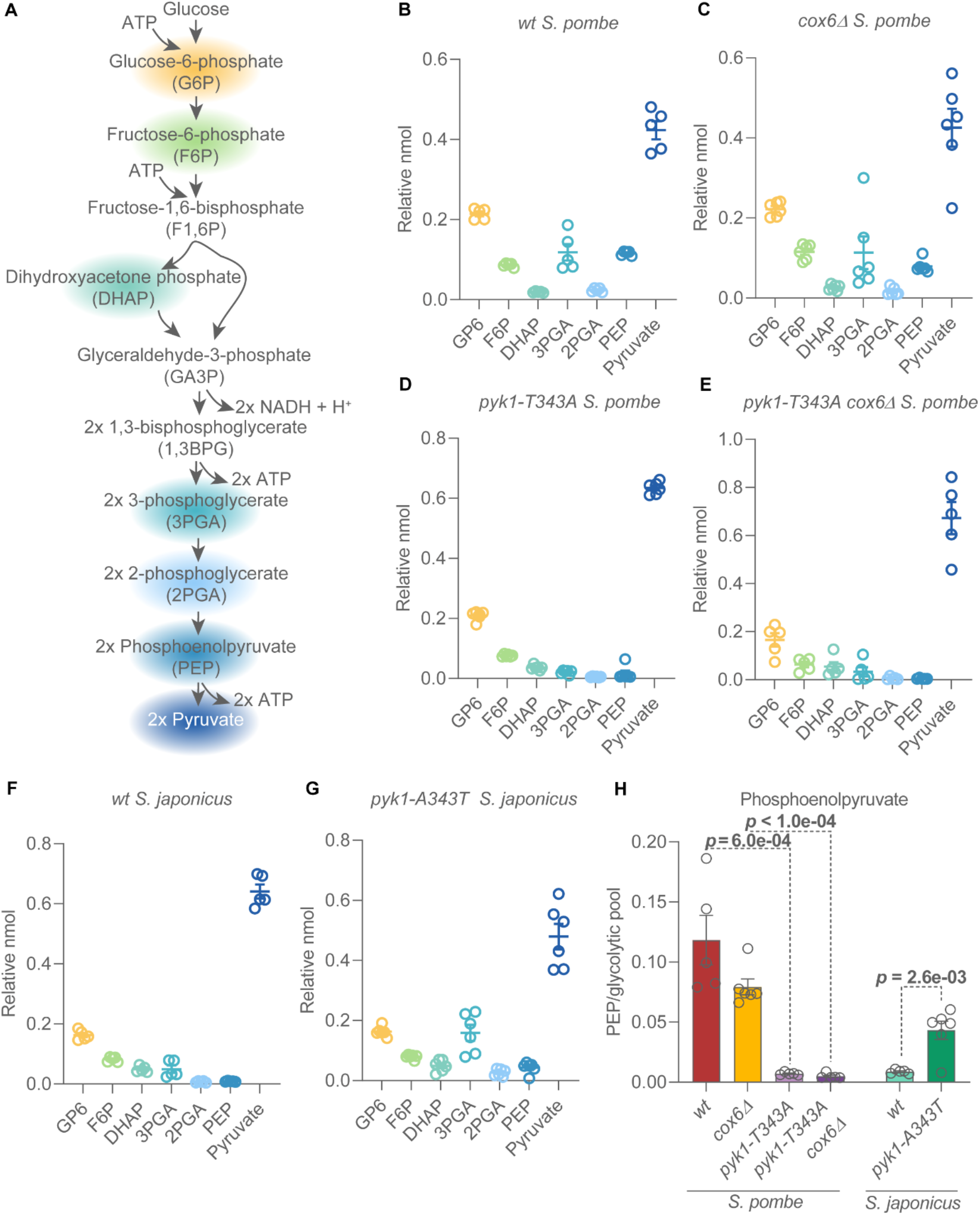
(**A**) Illustration of glycolysis, and the expected numbers of intermediate molecules per molecule of glucose catabolised, with quantified metabolites highlighted matching the colours used in the following plots. (**B**-**G**) Relative abundances of glycolytic intermediates in *S. pombe* and *S. japonicus*. Detected glycolytic intermediates were quantified and the relative proportion of each metabolite relative to the sum of the abundances of G6P, F6P, DHAP, 3PGA, 2PGA, PEP and pyruvate were plotted. (**H**) Phosphoenolpyruvate abundance relative to the sum of detected glycolytic intermediates (G6P, F6P, DHAP, 3PGA, 2PGA, PEP and pyruvate). (**B**-**H**) Plotted values are means ±SEM of three technical and two biological replicate experiments. *p* values were calculated using unpaired t-test.

**Supplemental figure 4, related to Figure 4.**
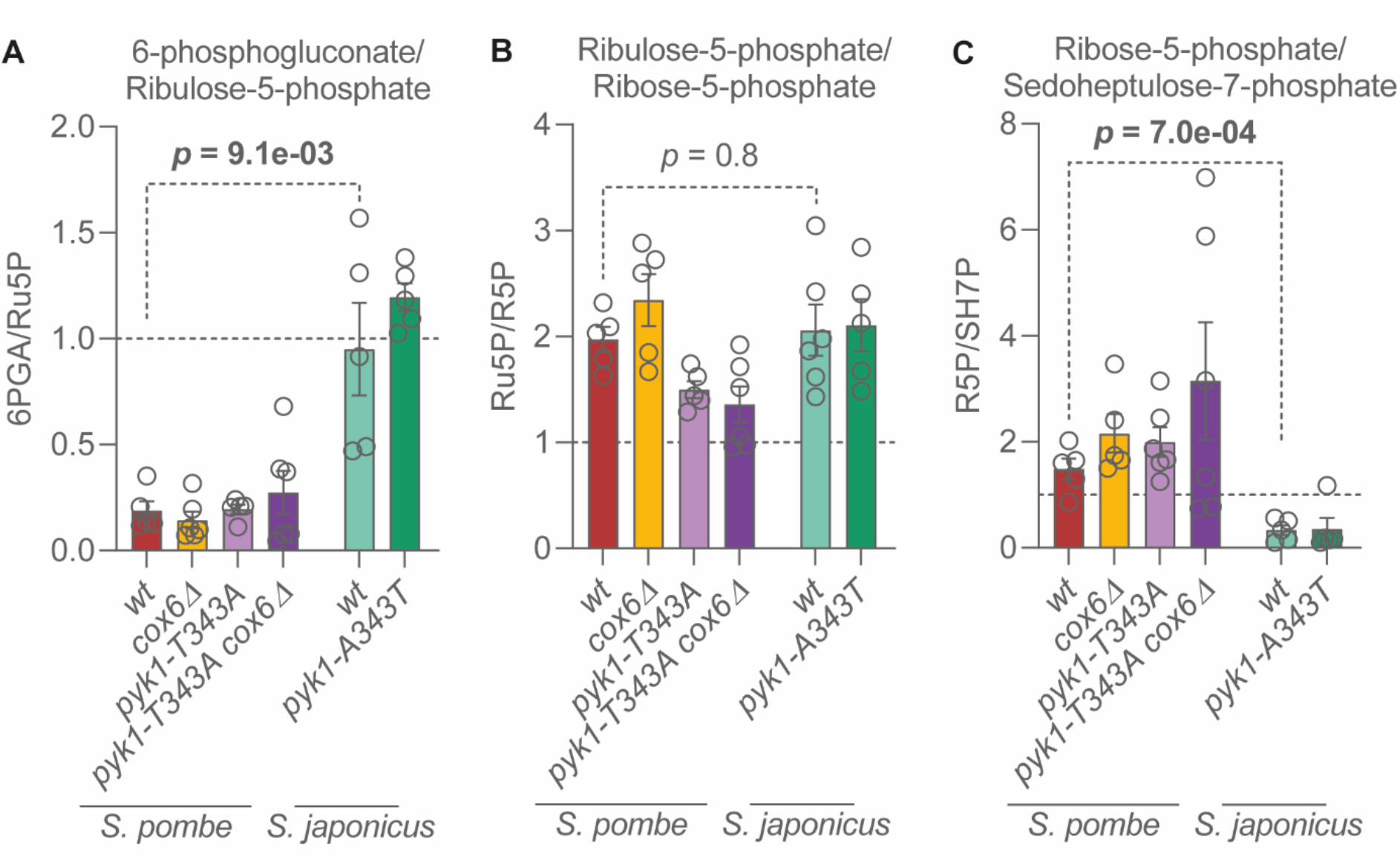
(**A**) Abundance of 6-phosphogluconate normalised to ribulose-5-phosphate. (**B**) Abundance of ribulose-5-phosphate normalised to that of ribose-5-phosphate. Ru5P appears to be more abundant than R5P, which may indicate a slow entry into the non-oxidative PPP in both *S. pombe* and *S. japonicus*. (**C**) Ribose-5-phosphate abundance normalised to sedoheptulose-7-phosphate. This ratio is below 1 in *S. japonicus* which indicates high levels of Sh7P. Sh7P synthesis also produces F6P, capable of re-entry into glycolysis. Plotted are the means ±SEM of two biological and three technical replicates. Dotted line indicates the ratio of 1. Statistical analyses were performed using unpaired t-tests.

## Notes

### Competing Interest Statement

The authors have declared no competing interest.

